# Early exercise disrupts a pro-repair extracellular matrix program during zebrafish fin regeneration

**DOI:** 10.1101/2024.11.15.623835

**Authors:** Victor M. Lewis, Rafael A. Fernandez, Samuel G. Horst, Carlos A. Gonzalez, Kryn Stankunas

**Affiliations:** Institute of Molecular Biology, University of Oregon, Eugene, Oregon USA; Department of Biology, University of Oregon, Eugene, Oregon USA

**Keywords:** Zebrafish, Regeneration, Hyaluronic acid, Yap, Mechanotransduction

## Abstract

Mechanical stimulation effects on cell behaviors that restore organ form and function during tissue repair are unresolved. We applied swim flume-mediated exercise during zebrafish caudal fin regeneration to explore mechanical loading impacts on a robust model of organ regeneration. Exercise initiated during but not after blastema establishment compromised fin regeneration, including outgrowth and skeletal pattern. Long-term tracking of fluorescently labeled fibroblasts showed exercise loading disrupted blastemal mesenchyme formation. Transcriptomic profiling and section staining indicated loading reduced an extracellular matrix (ECM) gene expression program, including for hyaluronic acid (HA) synthesis. As with exercise loading, HA synthesis inhibition or blastemal HA depletion impaired blastema formation. We considered if injury-upregulated HA establishes a pro-regenerative environment facilitating mechanotransduction. HA density across the blastema correlated with nuclear localization of the mechanotransducer Yes-associated protein (Yap). Exercise loading or HA depletion decreased nuclear Yap, and culturing primary fin fibroblasts on HA-coated surfaces induced Yap nuclear localization. We conclude early exercise during fin regeneration disrupts expression of an HA-rich ECM supporting Yap-promoted blastema expansion. These findings reveal that fin regeneration is acutely sensitive to the timing and intensity of mechanical loading, underscoring how biomechanical forces integrate with regenerative programs to guide robust tissue repair.

## Introduction

Biomechanical forces shape cellular behaviors during tissue regeneration by influencing cell state transitions, proliferation, and extracellular matrix (ECM) dynamics (1, 2). However, how mechanical stimuli interact with biochemical signaling and genetic programs to control such processes for robust regenerative outcomes is poorly understood. Investigating these interactions reveals how physical forces contribute to robust tissue repair and how mechanotransduction guides regenerative outcomes.

Zebrafish caudal fin regeneration offers a powerful model for studying the influence of biomechanical forces on regenerative mechanisms. Following amputation, zebrafish regenerate their fins—including the segmented bony ray skeleton—within 3 to 4 weeks (3). These appendages contain key cell types involved in mammalian skeletal repair, including osteoblasts, fibroblasts, and a stratified epidermis (3–5). While zebrafish fin regeneration is overtly spectacular, core processes such as cell dedifferentiation, blastema formation, and ECM remodeling are conserved across vertebrate models of tissue regeneration (6, 7). Therefore, zebrafish fin regeneration provides a compelling system for understanding how mechanical forces influence regenerative patterning.

Fin regeneration progresses through three key phases: acute wound repair, blastema formation, and outgrowth (3). Following epidermal wound healing, a regenerative blastemal mesenchyme forms by the distal migration of ‘activated’ intra-ray fibroblasts and wound-adjacent osteoblasts that de-differentiate to a pre-osteoblast (pOb) state(3, 4). Blastema establishment is complete by 3 days post-amputation (dpa). During establishment, a distal organizing center directs blastemal compartmentalization to form a stem-cell-like niche that balances proliferation and redifferentiation. The outgrowth phase then begins, during which mesenchymal pre-osteoblasts (pObs) remain localized to the lateral blastemal niche (8–10). Fibroblast-lineage niche cells secrete growth factors that maintain pObs in a proliferative state, while proximal osteoblasts provide redifferentiation cues (8). This balance of cell proliferation and orderly tissue maturation progressively restores the uninjured fin’s size, shape, and skeletal pattern.

Experimental studies that alter fin structure and assess hydrodynamic stress patterns indicate external shear stress and resulting internal tension regulate ray growth and branching, implicating mechanical loading in regenerative patterning(11). Yes-associated protein (Yap) functions as a mechanotransducer during regeneration by responding to cytoskeletal dynamics and mechanical forces to regulate cell proliferation, differentiation, and tissue growth (12, 13). The role of Yap signaling in regeneration is closely linked to cytoskeletal organization and mechanical inputs, with F-actin and cell density regulating Yap localization and activity(12). In regions of high cell density, Yap remains primarily cytoplasmic and inactive, correlating with cortical F-actin organization. In low-density regions, Yap is nuclear and active, aligning with diffuse F-actin structures (12). These findings suggest that mechanical forces transmitted through the cytoskeleton serve as upstream regulators of Yap signaling, directing regenerative outgrowth.

We use a swim flume system to investigate how controlled mechanical loading influences cell state transitions and signaling during fin regeneration (14). We find that exercise loading during blastema formation disrupts regeneration, with long-term live imaging of fluorescently marked fibroblast-lineage cells showing impaired blastemal mesenchyme formation. Transcriptomic profiling and *in situ* staining demonstrate that exercise loading suppresses hyaluronic acid (HA) synthesis. Experimentally depleting HA in non-loaded fish similarly disrupts blastema formation, further suggesting that HA establishes a pro-regenerative extracellular environment. To connect HA abundance with downstream cellular processes, we first show HA levels correlate with Yap nuclear localization in regenerating fins. Experimental manipulation through exercise loading or HA depletion reduces nuclear Yap and cell proliferation, while culturing primary fin blastemal fibroblasts on an HA-coated surface is sufficient to induce Yap nuclear localization. Our study reveals a stage-specific integration of extracellular matrix dynamics and mechanotransduction during fin regeneration, with potential relevance for strategies – including their timing – to promote tissue repair.

## Results

### Fin locomotion variably and differentially impact regenerative outcomes

To assess the impact of increased mechanical loading on zebrafish fin regeneration, we first defined the swimming ability of wild-type adult zebrafish to increasing water flow using a swim flume (*n*=12) (Fig. S1*A* and *B*)(15). We then implemented 30-minute swim exercise/day regimens at various fractions of the average maximum swimming velocity to caudal fin-resected fish, exercising fish from 1-3 days post amputation (dpa) (Fig. S1*C)*. Progressively increased swimming speeds correspondingly decreased the 3 dpa regenerated fin area (*n*=10) (Fig. 1*A*). Next, we explored any temporal biases for the negative effects of exercise on fin regeneration. We initiated 50% max speed exercise at different phases of the regenerative process and monitored regeneration throughout outgrowth (Fig. S1*D*). Zebrafish exercised beginning at 1 or 2 dpa showed reduced regenerated fin area through the 28-day course. In contrast, cohorts for which exercise commenced at 3 or 5 dpa ultimately regenerated as well as unexercised controls (*n*=6-10) (Fig. 1*B*; Fig. S1*E*). Therefore, daily exercise initiated prior to the outgrowth phase decreases regenerative capability. Notably, the early exercise regimen initiated at 1 dpa also produced skeletal patterning defects in the regenerated tissue, including the loss of branching morphogenesis of the most peripheral rays (83%, *n*=6), ray fusions between at least two lepidotrichia (67%, *n*=6), and a frequent failure of the principal peripheral rays to regenerate (83%, *n*=6) (Fig. S1*F-H*).

**Figure 1.**
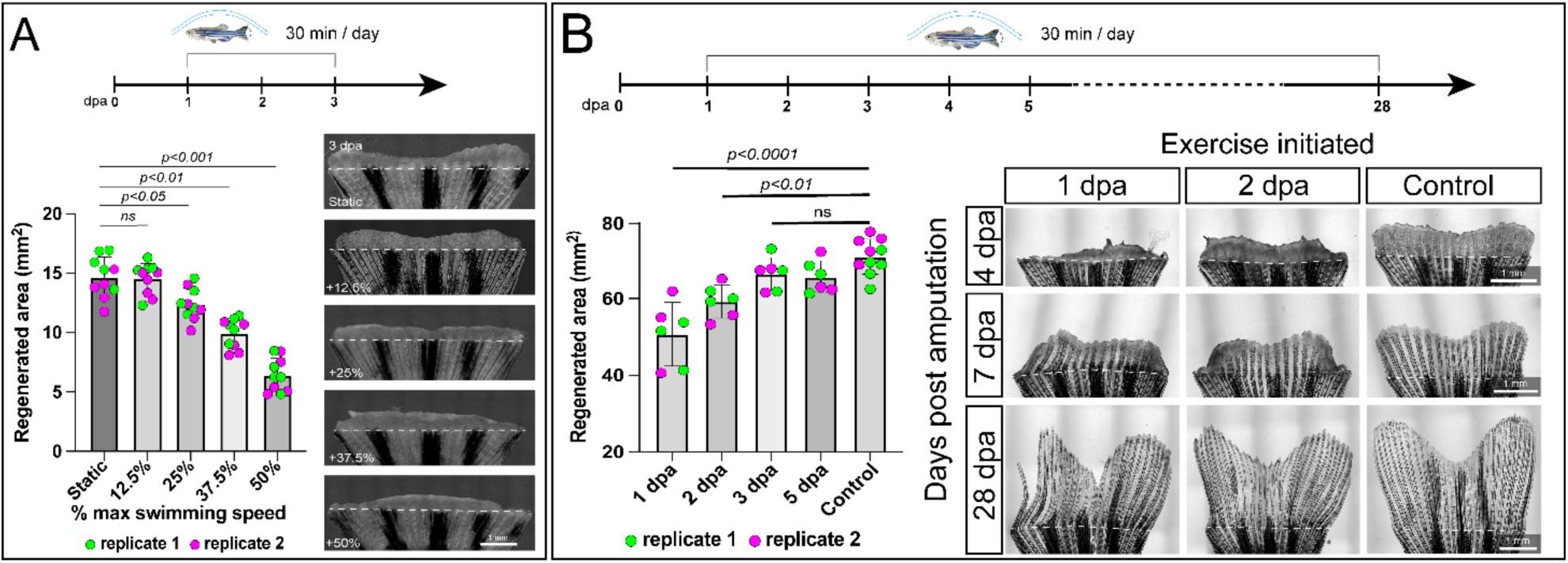
Early swimming exercise disrupts robust regeneration of the zebrafish caudal fin. (A) Quantification of regenerated area (mm^2^) at different percentages of maximum swimming speed (static, 12.5%, 25%, 37.5%, 50%) at 3 days post-amputation (dpa) (*n*= 5 fish/trial and 2 replicate trials). Representative whole-mount images are shown on the right for each swimming speed condition. (B) Quantification of regenerated area (mm^2^) at 28 dpa when exercise is initiated at different days post-amputation (1, 2, 3, and 5 dpa) and unexercised controls (*n*= minimum of 3 fish/trial and 2 replicate trials). Representative whole-mount images of fins at 4 dpa, 7 dpa, and 28 dpa are shown on the right for each exercise initiation condition. The timeline above each panel represents the exercise regimen (30 minutes/day) over the experimental period. White dashed lines show amputation plane. Scale bars represent 1 mm. Significance determined by one-way ANOVA with Dunnett’s post-hoc tests. Error bars represent standard deviation.

We next questioned if the modest swimming in standardized housing benefits regeneration. Preventing swimming by intubating and sedating caudal fin-resected fish for 16 hours starting at 2 dpa moderately reduced regenerated tissue (*n*=4) (Fig. S2) (16). We also considered if the adverse effects of exercise on regenerative capability might stem from systemic changes, such as alterations in metabolism or circulating factors, or localized effects within the caudal fin (17). To investigate this, we conducted swim trials after amputating both higher load-bearing caudal fins and lower load-bearing anal fins (18, 19). We observed a comparable loss of regenerated area of the caudal fin to previous experiments but only a modest reduction in anal fin regenerated area (*n*=6) (Fig. S3). The correlation between load intensity and extent of regeneration inhibition suggests localized effects within the recovering tissue rather than systemic influences. Taken together, these findings indicate that regular but non-intensive swimming produces local mechanical forces supporting robust caudal fin regeneration.

### Early swimming exercise inhibits blastema establishment

The observed decrease in regenerative potential when exercise was initiated at 1 or 2 days post-amputation (dpa) suggested that swimming exercise impedes blastema establishment. To explore this hypothesis, we employed longitudinal live imaging of fluorescently marked fibroblast lineage cells using the *Tg(sost:nlsEos)* line during the initial week of regeneration with and without swimming exercise (20, 21). Exercised cohorts showed a notable inhibition of fibroblast lineage establishment, with a more pronounced effect at the fin periphery (Fig. 2*A*). Blastemal fibroblasts robustly re-established following the cessation of daily exercise (*n*=6) (Fig. 2*B*). Therefore, exercise-induced loading actively restrains a fibroblast-lineage regenerative response.

**Figure 2.**
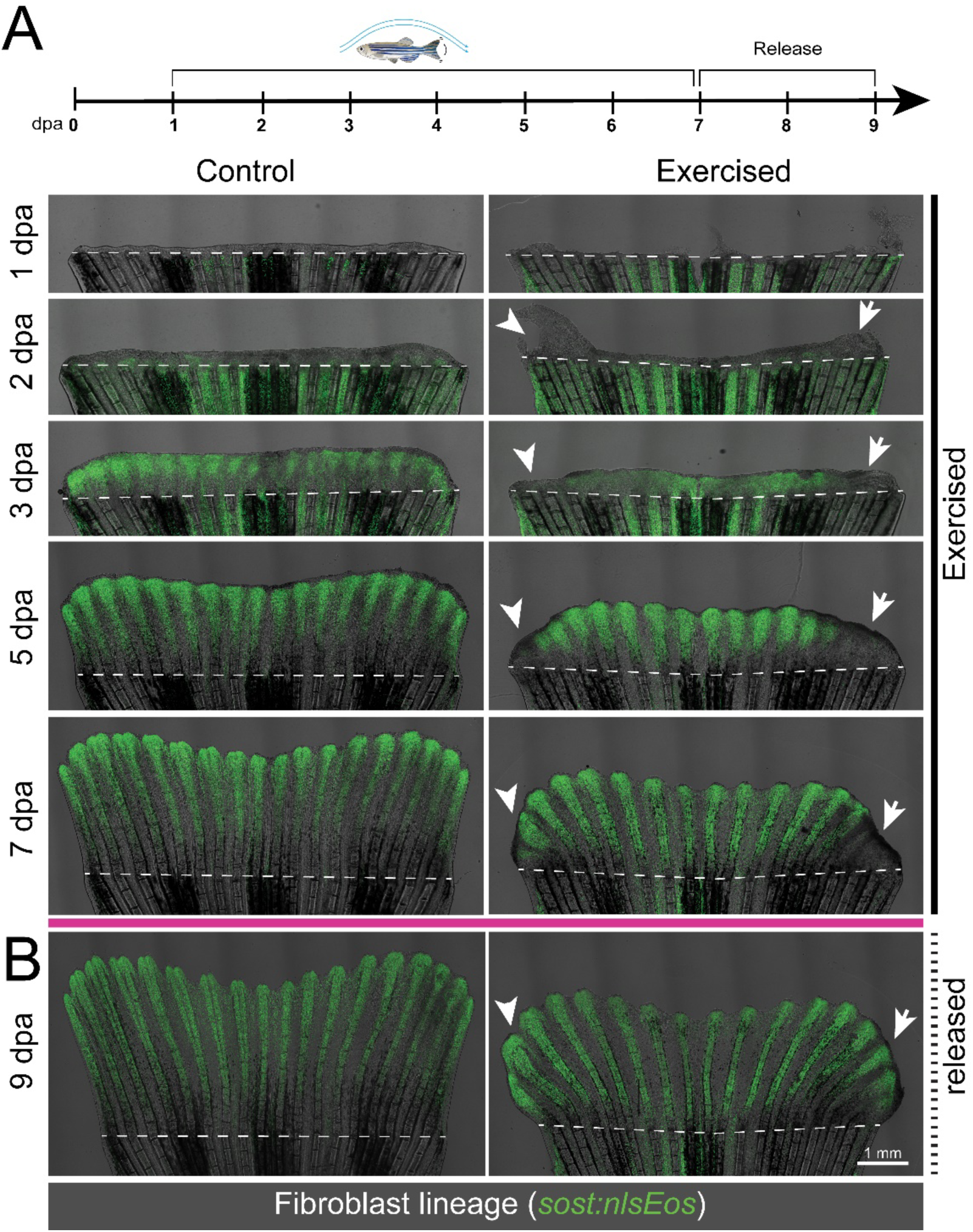
Early exercise initiation inhibits blastemal fibroblast establishment. (A) Representative 1-week time series of whole mount regenerating caudal fin overlay images in *Tg(sost:nlsEos)* blastemal fibroblast labeled animals (*n=6*). White arrowheads and arrows highlight either a delay or the complete loss of regeneration in peripheral regions, respectively, in the example shown. (B) Animal in (A) following release from exercise regimen showing robust re-establishment of fibroblast lineage cells. Dashed white lines indicate amputation plane. The scale bar shows 1 mm. The timeline above each panel represents the exercise regimen (30 minutes/day) over the experimental period.

We then employed fluorescent imaging of eosin-stained paraffin fin sections from animals exercised from 1-3 dpa to further characterize tissue-level changes in blastema establishment (*n*=3) (22). We observed a diminished blastema in sections of both central and peripheral rays, the separation of blastemal tissue from adjacent epidermis, and disorganization of the distal basal epidermis in sections from the more severely affected peripheral rays (Fig. S4). We conclude early exercise-induced loading during fin regeneration impairs wound epidermis organization and blastema establishment.

### Exercise during the blastema establishment phase disrupts ECM gene expression

To discern the molecular changes induced by exercise during regeneration, we performed transcriptional profiling (RNA-Seq) of caudal fin tissue from animals exercised from 2-3 dpa. We identified 124 significantly upregulated and 251 downregulated genes with levels changed more than 1.5x (Supplemental File 1). Examination of cell lineage-specific gene expression patterns supported the disruption of blastema-dependent regenerative events. Markers indicative of osteoblast maturation (*sp7, cdh11*) and vascular regeneration (*fli1a, fli1b*) were decreased, likely reflecting the dependence of these lineages on blastema-derived growth cues (Fig. S5*A*)(21, 23). Markers associated with fibroblast (*snai1a, snai1b, tph1b, tnfaip6*), pre-osteoblast (*runx2a, runx2b*), and basal epidermal lineages (*tp63, fras1*) remained largely unaltered whereas superficial epidermal markers (*krt4, epcam*) were increased (Fig. S5*A*)(21, 23). We assessed lineage-specific fluorescent reporters in swim-exercised fish to further validate the anticipated disruption in tissue distributions caused by swimming exercise. Consistent with our expectations, reporters marking immature and maturing osteoblasts [*Tg(sp7:gfp;runx2:mCherry)*], and vasculature [*Tg(fli1:gfp)*] all showed reduced contributions of their respective lineages after early exercise (*n*=6) (Fig. S5*B-E*)(20, 21, 24–26).

We next examined exercise-altered gene expression scores for signaling pathways implicated in fin regeneration. This analysis revealed a significant decrease only in components of Hedgehog/Smoothened (Hh/Smo) signaling (Fig. S5*F*). Consistently, expression of the *Tg(-2.4shha:gfpABC)^sb15^ sonic hedgehog a* reporter line was decreased in swim-exercised fish, including to negligible levels in the peripheral regions (*n*=6) (Fig. S5*G, H*)(25). As Shh signaling promotes branching morphogenesis, exercise-reduced *shha* pathway expression likely contributes to the observed peripheral ray branching defects (9, 27). However, chemical inhibition of Smo, which blocks detectable Hh/Smo activity, negligibly impacts fin outgrowth (9, 27). Therefore, while decreased *shha* activity likely contributes to exercise-disrupted peripheral ray branching morphogenesis, it is unlikely to account for the reduction in regenerative area.

We turned to an unbiased Gene Ontology (GO) of downregulated genes to identify potential processes underlying exercise-disrupted blastema establishment (Fig. 3*A*) (28). Extracellular matrix (ECM)-associated terms were the most highly over-represented terms (Fig. 3*A*), which predominantly reflected a common group of genes defining an exercise-disrupted ECM gene set (Fig. 3*B*). We investigated the lineage-specific expression of these ECM genes using our existing single-cell RNA-Seq (scRNA-Seq) dataset of regenerating fins, with a focus on distinct clusters representing superficial epidermis, basal epidermis, and combined blastemal fibroblast and osteoblast-lineage cells (Fig. 3*C*)(21). The ECM gene set was over-represented within the fibroblast and osteoblast lineage cluster, aligning with these lineages being most affected by exercise (Fig. 3*D*). Therefore, the ECM gene set comprises likely candidates accounting for swimming exercise disruption of blastema establishment.

**Figure 3.**
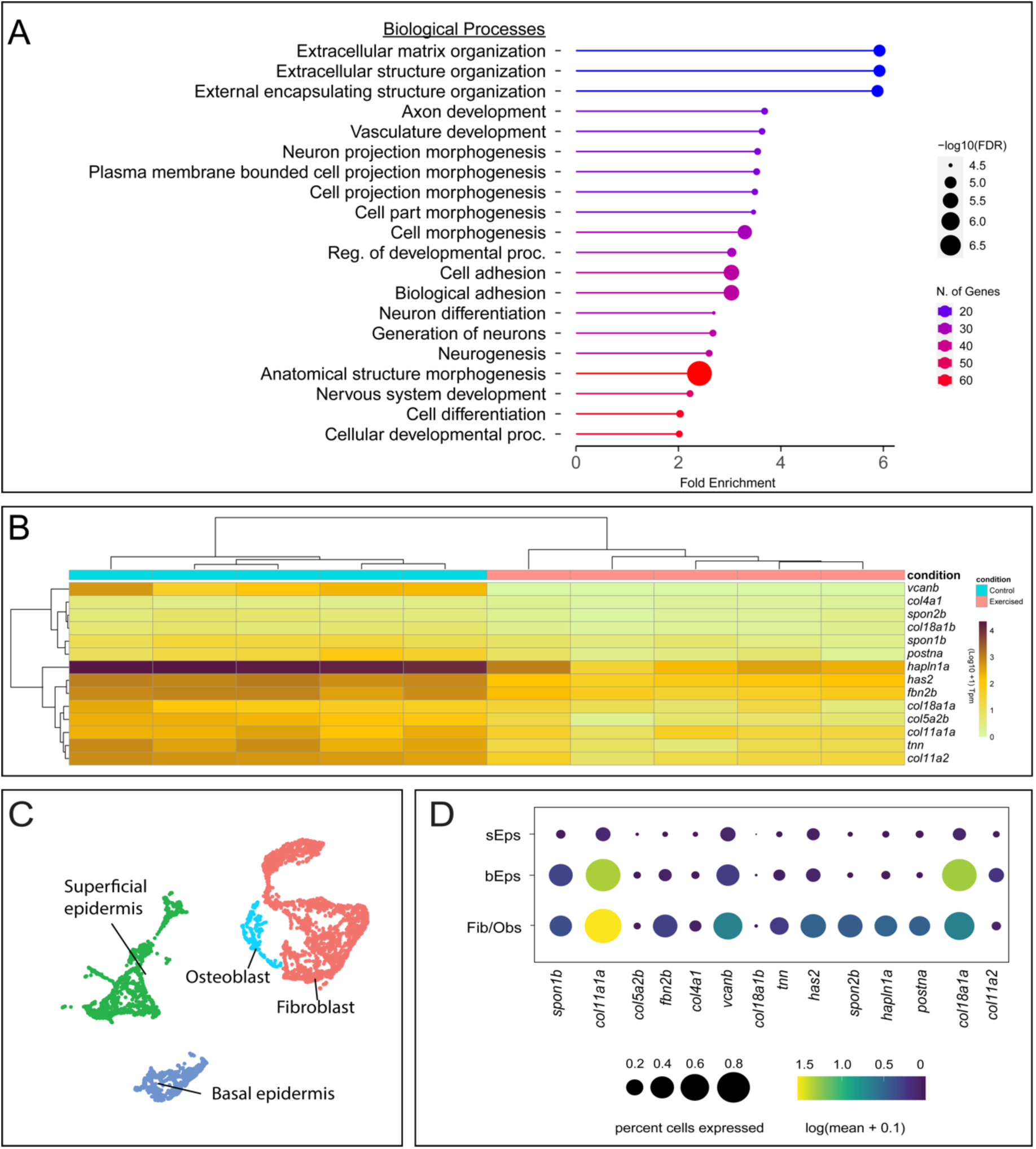
Transcriptomic profiling identifies an extracellular matrix (ECM) program downregulated in exercised animals. (A) Gene Ontogeny analysis showing the most significantly enriched biological processes from genes differentially downregulated in transcriptomic comparison. (B) Heatmap and hierarchical clustering of RNA-Seq replicates showing the expression profiles of genes identified within the top two ECM-related GO terms. (C) UMAP of previously published 3 and 7 dpa scRNA-Seq dataset, subset to clusters representing superficial epidermis, basal epidermis, and the combined blastemal fibroblast and osteoblast-lineage cells. (D) Bubble plot showing over-representation of genes identified within the top two ECM-related GO terms within the fibroblast and osteoblast lineage cluster.

### Early exercise decreases hyaluronic acid synthesis

We were intrigued by the presence of *hyaluronic acid synthase 2* (*has2*) within the ECM module given hyaluronic acid (HA) is a widespread ECM component that regulates cell proliferation and migration, including as a signaling ligand (29, 30). RNA-Seq transcript counts showed high expression of *has2* during caudal fin regeneration, lower but detectable expression of *has1*, and no *has3* expression, consistent with a previous report (Figure S6*A*)(29). *has1* and *has2* were significantly downregulated upon exercise in our RNA-Seq dataset, although *has1* failed to meet our fold-change threshold, potentially due to lower overall expression (Figure S6*B*). We next evaluated the presence of HA in regenerating tissues using biotin-conjugated hyaluronic acid binding protein (bHABP) staining of regenerating fin sections (31, 32). HA localized to the central blastema populated by fibroblast-lineage mesenchyme at 3 dpa (*n*=3) (Fig. 4*A*), matching previous reports, as well as the inter-and intra-ray connective tissue of unamputated regions (Fig. 4*B*)(31, 32). bHABP staining of 3 dpa cryosections from Tg(*sost:nlsEos*) regenerating fins showed high HA expression coinciding with endogenous Eos+ fibroblast lineage blastemal cells, and detectable but lower HA expression associated with Zn5-stained osteoblast-lineage blastemal cells (*n*=6) (Fig. 4*C*).

**Figure 4.**
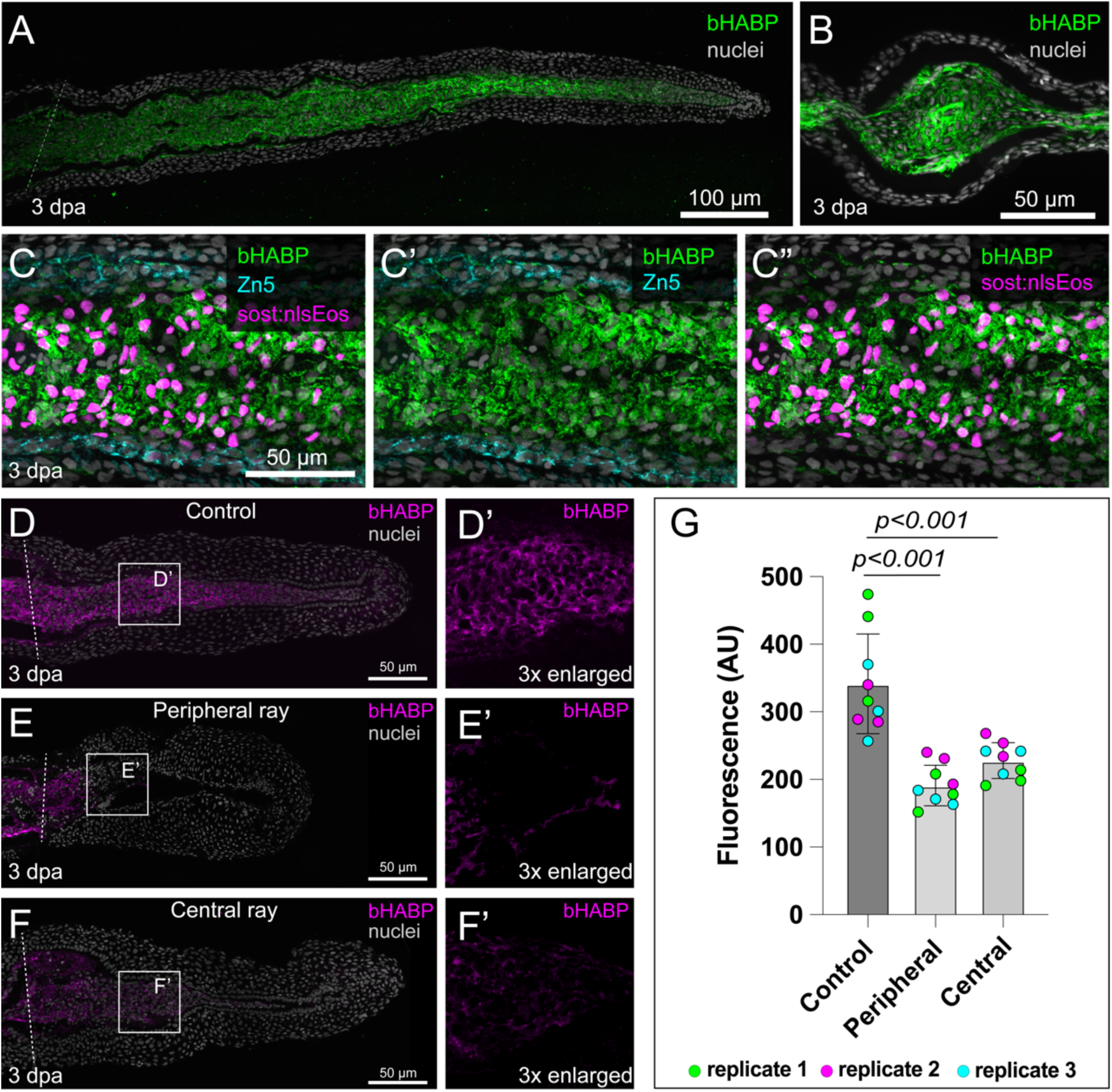
Exercise disrupts blastema-enriched hyaluronic acid production. (A) Representative confocal maximum intensity projection of a 3 dpa regenerating caudal fin paraffin section stained for bHABP (green) (*n*=3 sections; 1 animal). (B) Transverse paraffin section showing bHABP staining (green) within unamputated caudal fin tissue (*n*=3 sections; 1 animal). (C-C’’) Cryosection of a 3 dpa regenerating caudal fin from a Tg(*sost:nlsEos*) fibroblast lineage reporter fish showing co-staining of bHABP (green), Zn5 (cyan; osteoblasts), and Eos (magenta). Panels showing bHABP with Zn5 (C’) and bHABP with Eos (C’’) highlight cellular localization (*n*=6 sections; 2 animals). (D-F) Representative images of wildtype regenerating caudal fin paraffin sections at 3 dpa stained for bHABP (magenta) in control (D, D’) peripheral ray (E, E’) and central ray (F, F’) (*n*=9 sections; 3 animals). The boxed areas in panels D-F are shown 3x enlarged in panels D’-F’ to highlight bHABP staining. (G) Quantification of bHABP fluorescence intensity (AU) in control, peripheral ray, and central ray conditions from D-F (*n*= 3 sections/replicate and 3 replicate animals). Dashed white lines indicate the amputation plane. Scale bars represent 100 µm in panel A, 50 µm in panels B-F. Significance determined by one-way ANOVA with Dunnett’s post-hoc test. Error bars represent standard deviation.

bHABP staining of wildtype exercised animals’ regenerating fin sections after 2 days of swimming exercise (1-3 dpa) showed a significant reduction in HA, consistent with decreased *has1 and has2* transcript levels (*n*=9) (Fig. 4*D-G*). We tested if the reduced HA accounted for deleterious exercise effects by exogenously increasing HA in the regenerating blastema’s prior to exercise. To do so, we microinjected 10 nL of 0.625 ng/nL (low), 1.25 ng/nL (moderate) and 2.5 ng/nL (high) solutions of high molecular weight HA into individual blastemas of half the fin at 36 hours post amputation (hpa) followed by two swim exercise sessions at 48 and 72 hpa at speeds previously identified as deleterious (i.e. 25, 37.5 and 50%) (*n*=10) (Fig. S7*A-D*) (33). None of the HA treatments were beneficial at 50% speed. However, moderate HA addition (1.25 ng/nL) significantly improved outcomes relative to both mock and uninjected controls at 25 and 37.5% speeds (Fig. S7*D*, *E*). High HA concentrations (2.5 ng/nL) were deleterious at all speeds, likely due to the high viscosity of the injection solution (Fig. S7*D*, *E*). Nevertheless, the partial mitigation of exercise-dependent reductions in fin regeneration by moderate HA injection links the effects of early exercise to disrupted HA production during blastema establishment.

### Hyaluronic acid promotes blastema establishment and regeneration

We next explored the effects of chemically or enzymatically depleting HA independent of exercise. First, we validated in vivo use of 4-Methylumbelliferone (4-MU), a competitive inhibitor of hyaluronic acid synthesis (34), to reduce HA levels over a 0-3 dpa regeneration period (32) monitored by bHABP staining of regenerating fin sections. Adding 4-MU to fish water at concentrations from 0 to 300 μM dose-dependently inhibited HA deposition and decreased regenerated area (*n*=9) (Fig. S8*A, B*). We then used the *Tg(tph1b:mCherry)* fibroblast-lineage blastemal reporter (35) to show inhibiting HA synthesis through 150 μM 4-MU treatment from 0-3 dpa prevented activated fibroblasts from robustly populating the blastema, as seen with extensive exercise (*n*=12) (Fig. 5*A-D*).

**Figure 5.**
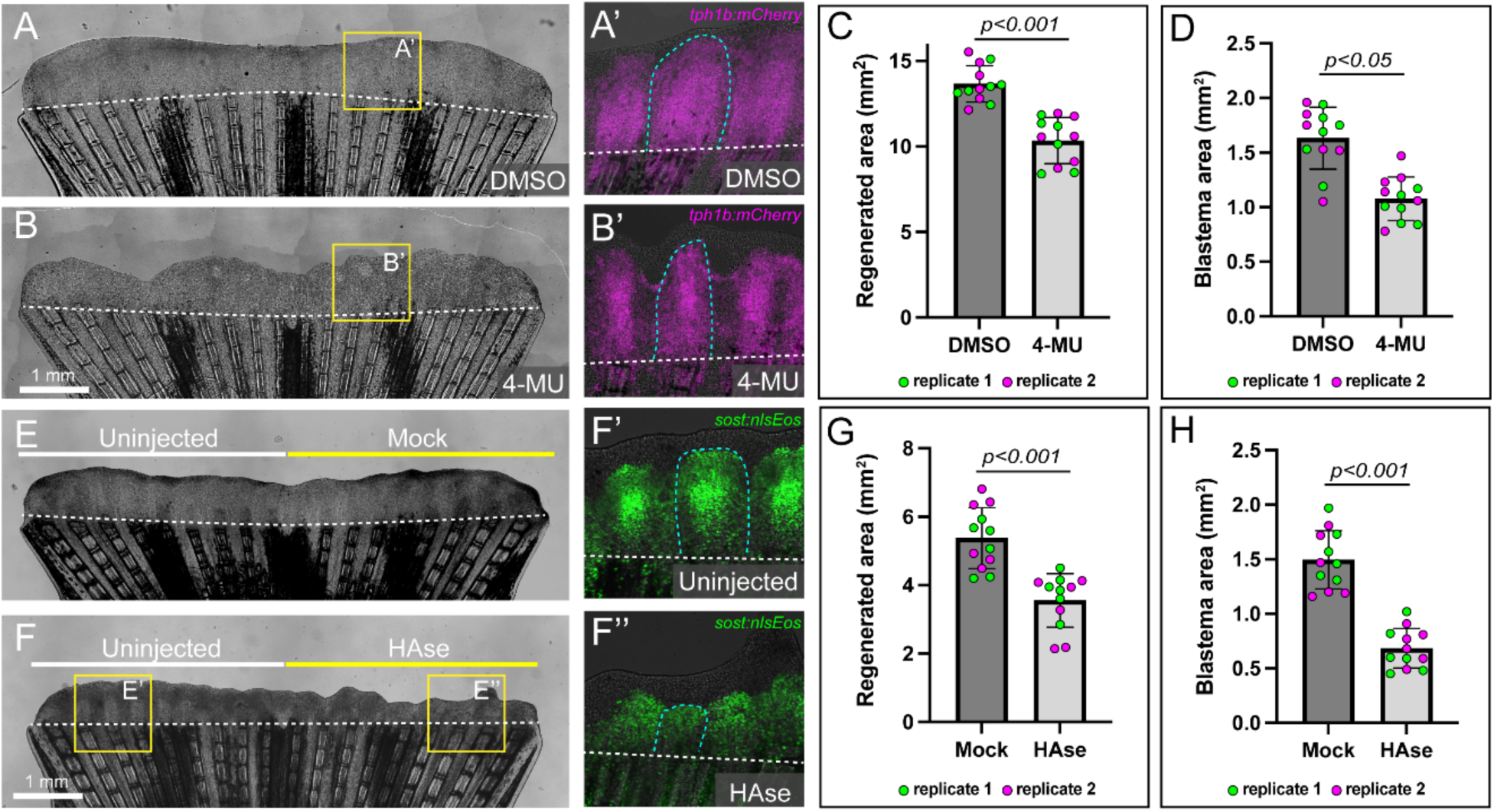
Hyaluronic acid supports robust blastema establishment during fin regeneration. (A, B) Representative regenerating caudal fins following 3 days treatment with 150 µM 4-MU (B) or DMSO controls (A) (*n=*6 animals/trial, 2 replicate trials). (A’, B’) Fluorescent overlay from A and B showing decreased fibroblast-lineage blastema establishment following 4-MU treatment in 3 dpa *Tg(tph1b:mCherry)* reporter animals. (C, D) Quantification of regenerated area (C) and blastema area (D) following 150 µM 4-MU from 0-3 dpa (E, F). Representative whole-mount fins after intra-blastemal injection of 15 mg/mL Hyaluronidase (HAse) (*n=*6 animals/trial, 2 replicate trials). (F’, F’’) Fluorescent overlay from F showing decreased fibroblast establishment relative to uninjected control tissue following blastemal HAse injection in 3 dpa *Tg(sost:nlsEos)* animals. (G,H) Quantification of regenerated (C) and blastema area (D) following HAse injection. Fluorescent lineage-reporter images are confocal maximum intensity projection overlays. Dashed white lines indicate amputation plane. Scale bars show 1 mm. Significance in (C, D) determined by unpaired two-tailed Student’s t-test. Significance in (G, H) determined by paired two-tailed Student’s t-test. Error bars represent standard deviation. Figure legend on next page.

As an additional assessment of HA-dependent blastema deposition, we depleted HA by 10 nL intra-blastemal injection of 15 mg/mL hyaluronidase during the establishment phase(36, 37) (Fig. S8*C*, *D*)(33). Hyaluronidase injection at 36 hpa decreased regenerative area and diminished blastema formation, as evidenced by reduced *Tg(sost:nlsEos)* expression at 3 dpa (*n*=12) (Fig. 5*E-H*). These chemical and enzymatic HA depletion experiments demonstrate that, independent of swimming exercise, injury-upregulated HA promotes blastema establishment and subsequent regeneration.

### Hyaluronic Acid mediates mechanotransduction-dependent proliferation through Hippo/YAP signaling

We sought to determine how exercise-sensitive HA production promotes robust blastema establishment. We first considered if HA could be acting as a signaling molecule affecting essential blastemal mesenchyme cell behaviors. However, we were unable to identify transcripts for the canonical HA co-receptors *cd44a/hmmr* in blastemal mesenchyme cells within our scRNA-Seq dataset (Fig. S9 *A, B*) (30, 38). Therefore, we considered if HA functions indirectly by enabling a pro-regenerative extracellular environment (39). The Yes-associated protein (Yap), a transcriptional co-activator of the Hippo signaling pathway, is required for fin regeneration and is linked to cytoskeletal interactions with the ECM (12). Yap responds to mechanical signals and plays a crucial role in regulating cellular behavior in relation to ECM stiffness (40, 41). Therefore, Yap likely functions as a mechanotransducer responsive to changes in the regenerative fin extracellular environment. We first confirmed previous reports of distal blastemal cytoplasmic Yap and nuclear localization in the proximal to medial blastema at 3 dpa (*n*=9) (12). We then combined bHABP and Yap staining of sectioned regenerating fins to correlate increased HA and Yap nuclear localization proximally and decreased HA and cytoplasmic Yap distally at 3 dpa (*n*=9) (Fig. S9*C-G*). These correlations suggest an HA-rich ECM supports blastemal establishment by enabling mechanosensitive Yap activity.

We tested whether HA promotes Yap-dependent mechanotransduction by monitoring Yap activity upon exercise-induced HA loss and experimentally induced HA depletion through chemical inhibition or enzymatic degradation (Fig. 6). Each scenario of depleted HA significantly reduced Yap nuclear localization (*n*=9) (Fig. 6*A-D*, *F, G*). Blastemal HA injections prior to swimming maintained nuclear Yap localization in exercised fish, further linking exercise effects to HA levels and downstream Yap activity (*n*=9) (Fig. 6*E*, *H*). Yap activity is necessary for blastemal proliferation during fin regeneration (12). Therefore, decreased Yap nuclear localization upon HA disruption could, at least in part, account for the deleterious effects of exercise on blastemal establishment.

**Figure 6.**
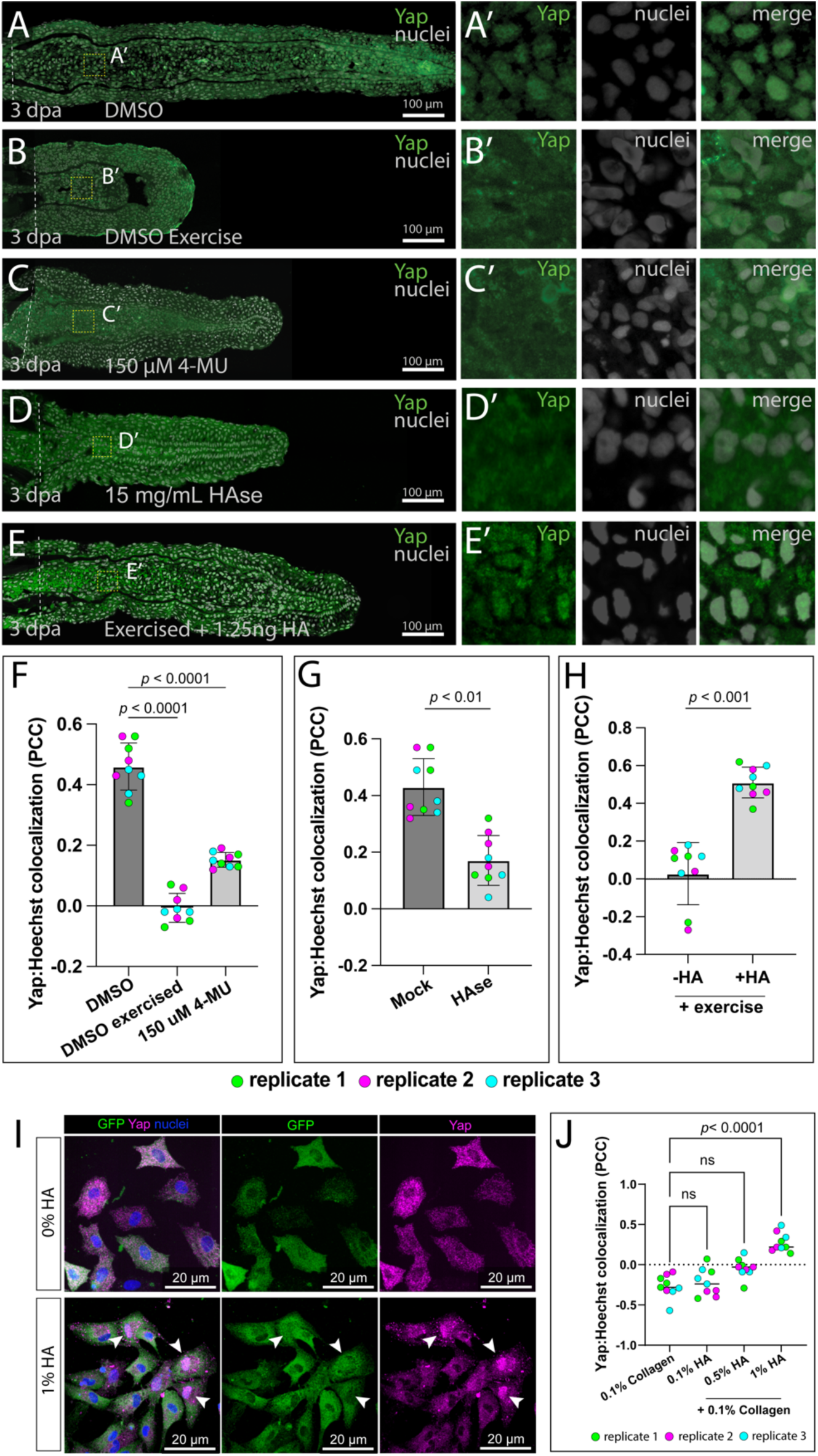
Hyaluronic acid supports Yap nuclear localization during fin regeneration. (A–E) Representative images of 3 days post amputation (dpa) regenerating fin sections stained for Yap (green) under different hyaluronic acid (HA)-altered conditions: (A) Unexercised DMSO controls. (B) Swim exercise for 3 days at 50% maximum swim speed. (C) 3-day treatment with 150 µM 4-methylumbelliferone (4-MU). (D) Intra-blastemal injection of 15 mg/mL hyaluronidase (HAse) at 60-hour post amputation. (E) Intra-blastemal injection of 1.25 ng/nL HA at 36 hpa followed by 2 swim exercise sessions at 50% maximum swim speed. The boxed areas in panels A-E indicate 3x enlarged in panels A’-E’ to highlight Yap colocalization with nuclei (n=3 sections; 3 animals). (F-H) Quantification of Yap-Hoechst colocalization (Pearson correlation coefficient, PCC) following (F) DMSO, exercised or 150 μM 4-MU treatment, (G) mock or intra-blastemal injection of 15 ng/nL HAse, and (H) with and without intra-blastemal injection of 1.25 ng/nL HA prior to exercise (*n*= 3 sections/replicate and 3 replicate animals). (I) Representative immunofluorescence images of primary fin blastema-derived fibroblasts cultured overnight on 0.1% collagen alone (0% HA) or supplemented with 1% hyaluronic acid (HA) (*n*=9 over 3 replicate trials). Cells were stained for GFP (green; marking *tph1b:GCaMP6s*⁺ fibroblasts), Yap (magenta), and nuclei (blue; Hoechst). Cells treated with 1% HA display enhanced nuclear Yap localization (arrowheads) (J) Quantification of Yap nuclear localization by Pearson’s correlation coefficient (PCC) of Yap and Hoechst signal across increasing HA concentrations (0%, 0.1%, 0.5%, and 1%) on 0.1% collagen. Each dot represents a single image; colored dots represent independent replicates. ns = not significant. Dashed white lines indicate amputation plane. Scale bars are as labeled. Significance in (F, J) determined by one-way ANOVA with Dunnett’s post-hoc test. Significance in (G, H) determined by paired two-tailed Student’s t-tests. Error bars show standard deviation.

We next tested whether exogenous hyaluronic acid (HA) is sufficient to induce Yap nuclear localization in regenerative fin fibroblasts. Primary cells were isolated from regenerating fins of Tg(*tph1b*:*GCaMP6s*) transgenic animals to selectively mark those of fibroblast lineage, and then seeded on a 0.1% collagen matrix (42). Co-immunostaining for endogenous Yap and GCaMP6s revealed that under basal conditions, Yap remained largely cytoplasmic, with minimal nuclear localization, independent of local cell densities across seeded culture plates (*n*=3) (Fig. S10*A*). Supplementation of the collagen coating with HA at increasing concentrations (0.1%, 0.5%, and 1.0% w/v) produced a concentration-dependent shift in Yap localization (*n*=9). At 0.1% HA, fibroblasts failed to show nuclear Yap enrichment, while 0.5% and 1.0% HA resulted in a subset of fibroblasts showing nuclear Yap enrichment (Fig. 6*F*; Fig. S10*B*). Quantification of the nuclear-to-cytoplasmic Yap fluorescence intensity ratios revealed a significant increase in nuclear Yap localization with 1.0% HA compared to collagen-only controls (Fig. 6*G*). These results demonstrate that exogenous HA is sufficient to drive Yap nuclear localization in primary fin blastema fibroblasts in vitro, consistent with a role for HA-dependent mechanotransduction in activating Yap signaling during regeneration.

Finally, we explored HA/Yap-dependent proliferation by evaluating Proliferative Cell Nuclear Antigen (PCNA) expression after both exercise-dependent and exercise-independent HA depletion. Exercise, chemical inhibition of HA synthesis, and enzymatic degradation of HA all significantly decreased the fraction of PCNA-positive nuclei (*n*=9) (Fig. S11*A-D*, *F, G*). Blastemal HA injections prior to exercise normalized PCNA-expressing nuclei counts (*n*=9) (Fig. S11*E*, *H*). Together, these findings indicate HA establishes a pro-regenerative extracellular environment enabling mechanotransduction-dependent cell proliferation during the establishment phase of fin regeneration.

## Discussion

We introduce the accessible zebrafish caudal fin regeneration model to study the impact of exercise induced mechanical loading on appendage and skeletal repair. We found that exercise loading early during fin regeneration impairs blastema formation and disrupts skeletal patterning, at least in part by reducing HA-enabled Yap activity. Exercise loading disrupts the normal expression of an ECM gene set that includes *has2*, *hapln1a* (hyaluronan and proteoglycan link protein 1), and several tissue repair-associated proteoglycans such as *vcanb* (Versican) and *postn* (Periostin)(43–46) Interestingly, HA and *hapln1a* are also involved in proper outgrowth and patterning during xenopus limb bud and mouse digit tip regeneration (29, 31, 32, 47). The ability of exogenous HA, added prior to exercise loading, to partially ameliorate the deleterious effects of early exercise further substantiates a rapidly growing body of evidence in support of HA‘s therapeutic potential (43, 48).

The reduced effect of exercise on the regeneration of the low-load-bearing anal fin compared to the high-load-bearing caudal fin suggests that the observed cellular and molecular impacts of exercise are likely caused by local, swimming-induced mechanical forces (18, 19, 49). However, we cannot entirely exclude the possibility that exercise produces systemic changes, such as elevated metabolic output. Future efforts to identify specific roles for mechano-sensors will be illuminating. Additionally, both overstimulated swimming activity and inhibition of movement through sedation decreased regenerative potential. It is important to note that the extent to which anesthesia itself might curtail regeneration remains unclear. Nonetheless, the similar deleterious effects of immobilization and overstimulated swimming suggest the existence of a mechano-transduction robustness network that ensures consistent blastema establishment across a range of “normal” swimming behaviors.

The most noticeable change in regenerated caudal fins subjected to exercise loading is a reduced regenerative capacity in the peripheral-most rays, while central rays exhibit a more moderate reduction in regeneration. This pattern aligns with previous studies suggesting that external shear stress and internalized tension are likely elevated at the dorsal and ventral margins of regenerating fins lacking a lobed structure(11). Furthermore, this may partially explain the diminished impact of exercise on regeneration observed after the transition to the outgrowth phase (∼3dpa), when the bilobed fin structure is re-established.

In addition to decreased regenerative capability, we observed that early exercise loading sensitizes regenerating fin skeletons to patterning disruptions. For example, the peripheral rays frequently failed to complete ray branching morphogenesis. Early exercise decreased expression of both basal epidermal *shha* and Hh pathway components, likely contributing to the skeletal branching defects given Hh/Smo signaling specifically promotes ray branching during fin outgrowth(9, 27). These observations align with previous reports suggesting that swimming mechanics influence both branching and *shha* activity (11). Future studies are warranted to determine how swimming-induced loading influences Shh signaling and to identify the mechanosensitive processes involved. Regardless, use of a swim tunnel for controlled swimming of regenerating fish will empower research into how exercise loading influences bone deposition and skeletal remodeling at different phases of the repair process.

Fibroblasts are vital for force transduction during skeletal rehabilitation exercises and are necessary for amphibian limb and mouse digit tip regeneration(6, 50–52). Likewise, zebrafish studies centrally implicate fibroblast-lineage state transitions and cell behaviors in fin regeneration (10, 21, 53–55). Both our transcriptomic analyses and lineage-specific transgenic reporter assays primarily identified fibroblasts and fibroblast lineage-dependent processes in the response to exercise during regeneration. Our analysis of genes downregulated by swimming exercise highlighted extracellular matrix changes underlying deleterious effects of exercise. It will be interesting to explore if the ECM gene set represents a co-regulated module in fibroblasts during regeneration and other repair contexts, including examples of mammalian skeletal injuries. Additionally, investigation into the genes upregulated by swimming exercise could provide further insights linking beneficial exercise effects observed in other studies with fibroblast lineage cellular processes (56).

ECM components play a crucial role in tissue homeostasis and signal transduction, particularly during the mechano-biological response to loading(57–60). Our study implicates fibroblast-produced ECM in creating a pro-regenerative environment essential for fin appendage regeneration. Specifically, we elaborate on prior reports that hyaluronic acid (HA) enables robust regeneration(29, 48), presenting evidence that HA facilitates blastema formation during the early establishment phase. Further, blastemal HA facilitates activity of the mechanotransducer Yap. Previous studies indicate Yap activity is initiated shortly after injury and required for blastema formation (12, 13). An intriguing model is that HA influences tissue stiffness in the fin environment, as shown in both hydrogel models and pancreatic islet inflammation *in vivo* (61–63). This encourages future investigations into ECM influences on cell-autonomous responses during regeneration or wound healing.

The interplay between mechanical stimuli and cellular reactions modulates regenerative processes, notably for bone remodeling and repair (64, 65). Cells and tissues possess a remarkable ability to sense and translate mechanical signals into essential biochemical cues, underscoring the importance of rehabilitative exercises (66, 67). This dynamic interaction has helped justify early rehabilitation even while other studies suggest premature rehabilitative exercise disrupts initial healing and thereby worsens outcomes following skeletal injuries (68). The latter view aligns with our observation of detrimental effects of exercise loading on fin regeneration while the blastema is being established but not after outgrowth initiates. Similar results were found in rat femur repair, where early mechanical loading proves detrimental, while delayed loading improves healing (68). Likewise, rehabilitative-like exercise shown here and in mammals appears beneficial up to a certain threshold, beyond which deleterious effects occur (69, 70). Together, our observations suggest fin regeneration shows similarities to the response of mammalian injuries to rehabilitative exercise. Whether these similar outcomes represent conserved underlying cellular and molecular processes or convergence on a beneficial repair paradigm remains unclear. Regardless, the phenotypic similarities underscore the value of using zebrafish fin regeneration to uncover the mechanisms by which mechanical loading influences skeletal appendage repair.

## Materials and Methods

### Zebrafish husbandry, fin amputations and regenerative area analysis

Zebrafish age 3-12 months were housed at approximately 28.5°C. Animals were maintained on a 14:10 day:night cycle and fed twice daily. The study used established zebrafish lines: wildtype (ABC), *Tg(sp7:EGFP)^b1212^*(26), *Tg(RUNX2:mCherry)*(9), *Tg(tph1b:mCherry)*(35), *Tg(sost:nlsEos)* (20, 21), *Tg(fli1:EGFP)*(24)*, Tg(krt4:dsRed)*(71), and *Tg(shha-2.3ABC:EGFP)*(25). Transgenic lines were maintained on the ABC strain background. Caudal or anal fins were amputated with a razor blade and then fish were returned to circulating fish water. Fish were euthanized by MS222 overdose. All zebrafish procedures were approved and monitored by the University of Oregon Institutional Animal Care and Use Committee (IACUC), adhering to the guidelines and recommendations outlined in the Guide for the Care and Use of Laboratory Animals (National Academic Press).

### Zebrafish swim flume calibration, swim fitness testing and exercise regimen

A 90 L swim tunnel (Loligo SW10275) was calibrated for water velocity by induction using the provided water vane velocimeter. Swimming fitness tests followed established critical swimming speed (uCrit) protocols with initial increases of 5 cm/s in velocity every minute for 5 minutes, followed by increases of one cm/s in velocity every minute thereafter until failure. For exercise and regeneration experiments, fin-resected zebrafish were allowed 24 hours recovery before swimming was initiated at times and speeds indicated. Fish were returned to normal flow on racks in the centralized aquatics facility after all swimming sessions.

### Long-term live sedation and imaging

Fish were anesthetized with 168 mg/L buffered tricaine and intubated in a 3D-printed chamber equipped with a tubing system modified from Castranova et al. (16). Intubated fish were submerged in water inside a walled coverslip (Cellvis Chambered Coverglass System #1.5 High Performance Cover Glass) and held down by thin strips of medical gauze placed over their trunk. Warmed and oxygenated water (using an air stone; Pawfly MA-60) containing 126 mg/L tricaine was delivered continuously using peristaltic pumps (Isco Tris 3-Channel Peristaltic Pump). The water supply was maintained between 24 and 28°C using a hot plate set to 35°C with continuous stirring (Corning PC-4200). Fish were kept intubated and sedated on the system for 16 hours. The following morning, fish health was confirmed by observing blood flow in the caudal fin and a steady heartbeat. Fish were revived by switching to regular system water while still intubated until independent breathing and movement was observed, at which point fish were returned to tanks. All fish restored normal swimming behavior.

### Imaging

Images were captured using a Nikon Eclipse Ti-E widefield microscope equipped with a Yokogawa CSU-W1 spinning disk confocal scanner unit or a Zeiss LSM 880 laser scanning confocal microscope. Confocal image stacks were processed using ImageJ software to produce single optical slice digital sections and/or maximum intensity projections. Adobe Photoshop was employed to adjust display levels, with consistent processing settings applied within each experiment.

## Statistical analysis

Fin morphometrics and section staining quantifications used ImageJ-Fiji (NIH) with data tables then organized with Microsoft Excel. RNA-Seq statistical analyses were conducted with Sleuth in Python. All other statistical analyses were conducted, and graphs created using GraphPad Prism v9.

### Paraffin section immunostaining and analysis

Amputated caudal fins were fixed in 4% paraformaldehyde (PFA) in PBS overnight at 4°C. The fins then were processed for paraffin sectioning, sectioned, and stained following established procedures (8). After washing with PBS, fins were decalcified for 4 days in 0.5 M EDTA, pH 8.0, with daily solution changes. Dehydration was achieved through an ethanol series, followed by clearing with xylenes, and embedding in paraffin wax. 7 mm sections were obtained using a Leica RM255 microtome. Antigen retrieval on rehydrated sections was performed using 1 mM EDTA with 0.1% Tween-20 (PCNA; SigmaAldrich) or 0.05% citraconic anhydride (Yap; Santa Cruz Biotechnology [sc-15407]) for 5 minutes in a pressure cooker. Following PBS washes, sections were blocked in 1x PBS with 10% nonfat dry milk for a minimum of 1 hour. Primary antibodies were diluted in blocking solution and applied overnight at 4°C. Sections were washed in PBS containing 500 mM NaCl with 0.1% Tween-20. Alexa Fluor conjugated secondary antibodies (Thermo Fisher) were diluted 1:1000 in blocking buffer and applied to sections for 1 hour at room temperature. After washing, nuclei were labeled with a 1:1500 Hoechst dilution (stock at 10 mg/mL) and the slides were mounted using SlowFade Gold Antifade (Thermo Fisher). The ImageJ-Fiji coloc2 tool was used to determine Yap:Hoechst co-localization. HA/Yap correlation was conducted in R using the cor function and ‘pearson’ method. ImageJ-Fiji cell counter tool was used to quantify PCNA+ cells. All images are confocal maximum intensity projections unless otherwise specified.

## Hyaluronic acid labeling and analysis

Hyaluronic acid (HA) staining used a biotinylated Hyaluronic Acid Binding Protein (bHABP; MilliporeSigma [385911]) at 1:100 dilution applied as described for immunostaining without antigen retrieval or NaCl wash and using a streptavidin-Alexa Fluor 647 conjugate secondary (Thermo Fisher) at 1:750 dilution. The ImageJ-Fiji measure tool was utilized to determine bHABP relative fluorescence intensities (RFU). All images are confocal maximum intensity projections.

### *In vivo* hyaluronic acid synthesis inhibition

4-Methylumbelliferone (4-MU, Sigma Aldrich) was dissolved in DMSO to establish a 150 mM stock then stored at-20°C. Immediately before use, 4-MU was thawed and further diluted in fish water. Animals were treated with 4-MU daily for 8 hours during the treatment window and then returned to system water.

### Intra-blastemal injections of Hyaluronic Acid and Hyaluronidase

Intra-blastemal injections were as previously performed(33). Briefly, fish were placed on their side on a solid agarose plate under a dissecting microscope. 15 mg/mL Hyaluronidase (Sigma #H3884) or Hyaluronic acid (1.3 - 1.8 MDa; Sigma #53747) was injected into each blastema using a glass needle and a micro-injection rig. For HA injections, half of the fin was injected with HA plus buffered phenol-red co-injection dye and the other half left uninjected, alternating fish between dorsal or ventral injections. For Hyaluronidase experiments, half of the fin was injected with Hyaluronidase plus co-injection dye and half with co-injection dye alone. Following injection, fish were returned to water and allowed to recover for at least 12 hours prior to swimming exercise or imaging.

### RNA-Seq library construction, sequencing and analysis

Fish with caudal fin amputations were exercised twice at 48 and 72 hours post amputation and then whole regenerate fin tissue collected into Trizol reagent on ice 4 hours later. RNA was prepared using a Direct-Zol RNA prep kit (Zymogenetics) and eluted into water. RNA was pooled from 2 male and 2 female size-matched fish for each replicate with replicates spanning two independent swimming cohorts. RNA was assessed and quantified using a nanodrop and fragment analyzer prior to library construction. Libraries were prepared with 1 μg of input RNA using a TruSeq library prep kit, indexed and pooled, and sequenced using an Illumina NovaSeq 6000 to a read depth of approximately 25 million reads per library. Reads were aligned with Kallisto (Pachter lab) against zebrafish reference transcriptome (GRCz11). Downstream analysis, including differential gene expression and graphics were conducted in Sleuth (Pachter lab)(72, 73). Gene Ontogeny analysis for biological processes used downregulated DEGs (*q<*0.05 and β>|1.5|) input into ShinyGO 0.77 with the zebrafish reference transcriptome, an FDR cutoff of 0.05 and 20 pathways shown with otherwise default settings(28).

### Single cell RNA-Seq analysis

For single cell RNA-Seq expression profiling, we used a pre-established and validated dataset prepared from regenerating fin tissue collected at 3-and 7-days post amputation(21). To focus on specific cell lineages, the “choose_cells” function was employed to restrict the extended dataset to exclusively epidermal, fibroblast, and osteoblast lineages. Uniform Manifold Approximation and Projection (UMAP) plot illustrating cell-type clustering and bubble plots showing cluster-specific candidate gene expression profiles was generated with R using Monocle3(74), the “Plot_cells” and “Plot_bubble” functions, respectively, with default parameters for all other settings.

### Primary Cell Isolation and Staining

Primary cells were isolated from regenerating caudal fins of *Tg(tph1b:GCaMP6s)* (42) zebrafish at 4 days post-amputation (dpa), following an adapted protocol from Stewart et al., 2014 (75). Excised fin tissue was surface-disinfected in 10% bleach, finely minced using a sterile razor blade, and enzymatically dissociated with Liberase DH (Sigma-Aldrich) to generate a single-cell suspension. Cells were plated onto glass coverslips coated with 0.1% collagen and varying concentrations of hyaluronic acid (Sigma-Aldrich) and incubated overnight at 30°C in an ambient air incubator. Cultures were maintained in Leibovitz’s L-15 medium (Thermo Fisher Scientific) supplemented with 20% fetal bovine serum (FBS; Sigma-Aldrich), 1% Penicillin-Streptomycin, and 1% GlutaMAX (Thermo Fisher Scientific). Primary cell cultures were stained using the same immunostaining protocol as for paraffin sections, omitting antigen retrieval. The Yap primary antibody (Santa Cruz Biotechnology, sc-101199) was used at a 1:200 dilution, and the GFP (GCaMP) primary antibody (Aves, GFP-1020) at 1:1000. Nuclear co-localization of Yap with Hoechst was quantified using the Coloc2 plugin in ImageJ/Fiji.

## Data Availability

RNA-Seq data has been deposited to the NCBI Gene Expression Omnibus, accession GSE293062. Requests for materials should be addressed to K.S.

## Acknowledgments

We thank the University of Oregon AqACS Facility for zebrafish care; the University of Oregon zebrafish community for support; Judith Eisen for reagents; Astra Henner for technical assistance and the Stankunas lab for input. The National Institutes of Health (NIH) provided research funding (R01GM149999). We acknowledge support by the Wu Tsai Human Performance Alliance and the Joe and Clara Tsai Foundation. S.G.H. and C.A.G. were supported by the University of Oregon Developmental Biology Training Program (T32HD007348). V.M.L. and S.G.H. were funded by NIH NRSA fellowships (F32GM140712, and F31HD113401, respectively).

## Notes

### Competing Interest Statement

The authors have declared no competing interest.

### Summary of Updates

Revised abstract; Introduction updated with additional background information; New line of experimentation using primary regenerating fin cells; Figures 3, 6, S4 and S10 revised with associated changes to main text; New Figure S11; Methods updated to reflect new experiments; Authorship, contributions, and funding updated.

