## Supplemental Figures for "Early exercise disrupts a pro-repair extracellular matrix program during zebrafish fin regeneration"

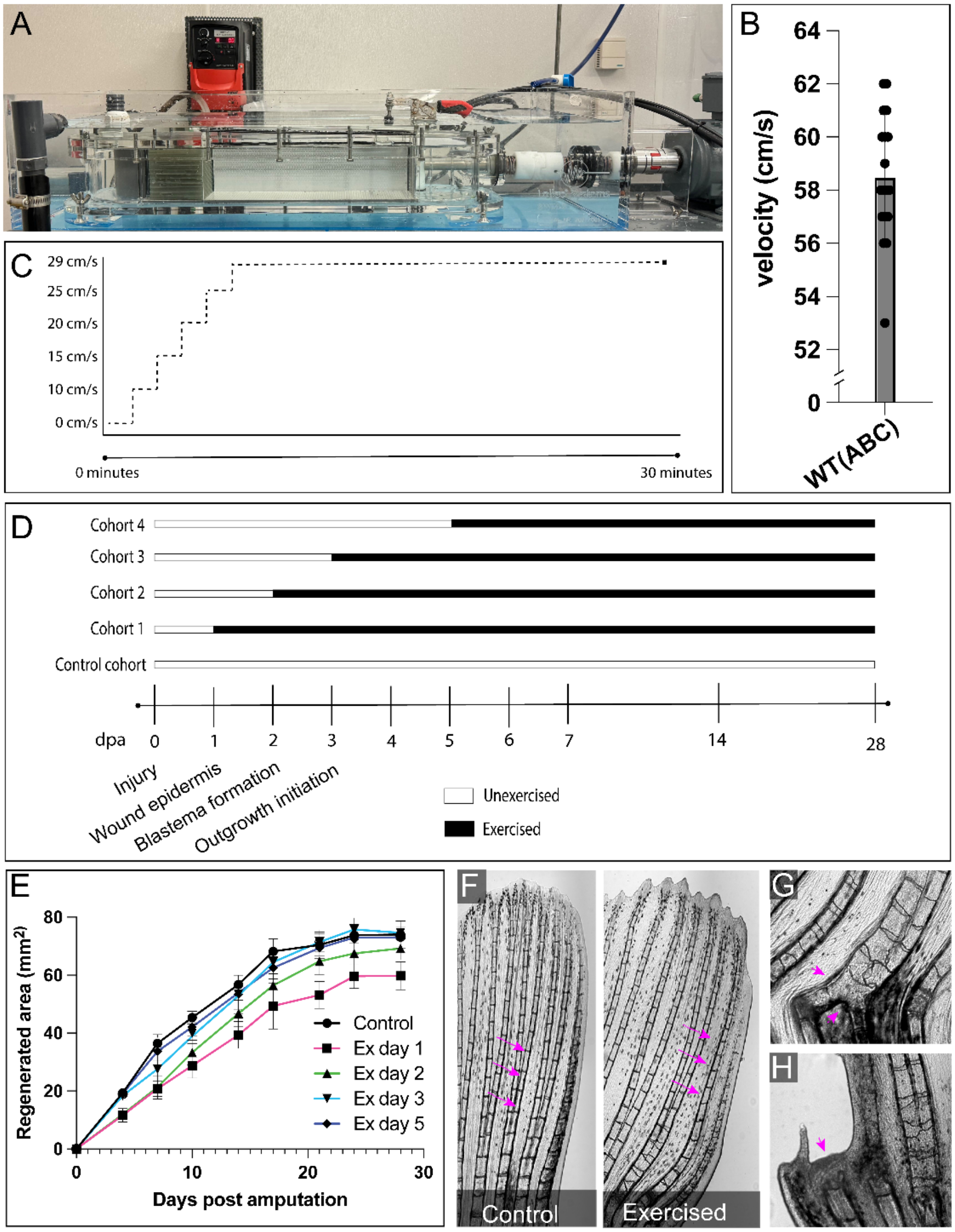


**Supplemental Figure 1. A controlled swimming approach to assess temporal effects of exercise on fin regeneration growth and skeletal patterning.** (A) Loligo swim flume used in experiments. (B) Graph showing the maximum swimming speed of wildtype zebrafish (*n*=12). (C) Schematic representation of daily exercise regimen. (D) Schematic detailing experimental design for delayed exercise onset cohorts. (E) Graph showing regenerated fin area over a full course of regeneration for the cohorts outlined in (D) (*n*=6/group). (F) Fin regions showing the loss of skeletal branching in peripheral rays. (G) Bony ray fusions following swimming exercise. (H) Inhibition of regeneration in principal peripheral rays. Magenta arrows indicate regions of interest. Dashed white lines indicate the amputation plane. Error bars represent standard deviation.

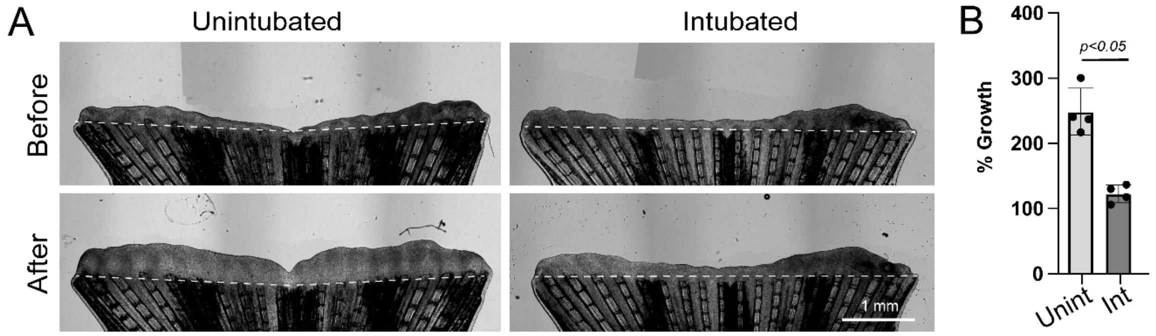


**Supplemental Figure 2. Swimming inhibition by anesthesia impairs fin outgrowth.** (A) Representative whole mount images of regenerating caudal fins before (48 hours post amputation [hpa]) and after (64 hpa) long-term sedated intubation. (B) Percent regenerative growth following inhibition of swimming activity through long-term anesthesia (*n*= 4 paired animals from individual trails). Unint = unintubated, Int = intubated. Dashed white lines indicate the amputation plane. Scale bars show 1 mm. Significance determined by a paired two-tailed Student's t-test. Error bars represent one standard deviation.

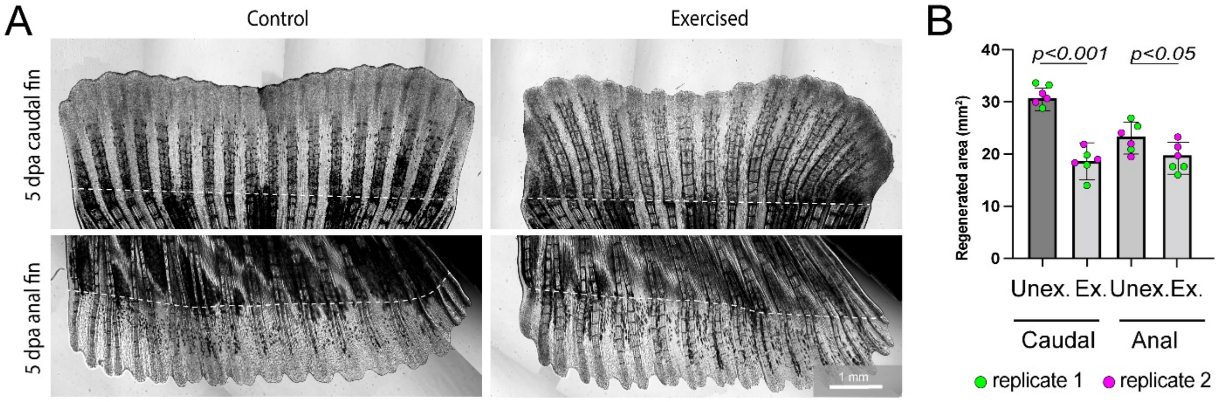


**Supplemental Figure 3. Exercise in high and low-load bearing fins suggests fin-autonomous swimming effects.** (A) Representative images of high-load bearing caudal and low-load bearing anal fins at 5 dpa following daily exercise (n=6). (B) Regenerated area in caudal and anal fins at 5 dpa following daily exercise (*n*= 3 animals/trial and 2 replicate trials). Unex = unexercised; Ex = exercised. Dashed white lines indicate the amputation plane. Scale bars show 1 mm. Significance determined by paired two-tailed Student's t-tests. Error bars represent standard deviation.

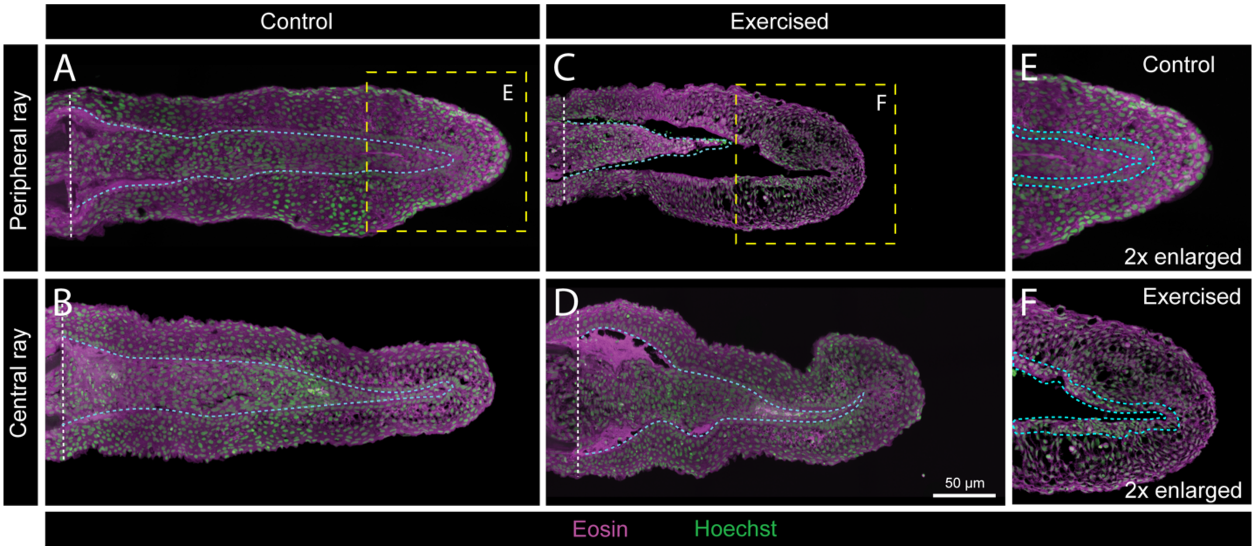


**Supplemental Figure 4. Early exercise disrupts wound epidermis organization and blastema integrity in regenerating caudal fins.** (A-D) Representative 3 dpa regenerating caudal fin sections from control (A, B) or exercised (C-D) fish stained with Eosin (magenta) and Hoechst (green). Sections show either peripheral (A, C) or central (B, D) rays following 3 days of exercise at 50% maximum swim speed. (E, F) 2x enlarged view of the yellow boxed areas indicated in panels A and C, respectively, showing detailed tissue organization of distal fin regions. Cyan dashed lines in E and F highlight the basal epidermis. White dashed lines show amputation plane. Scale bar shows 50 µm.

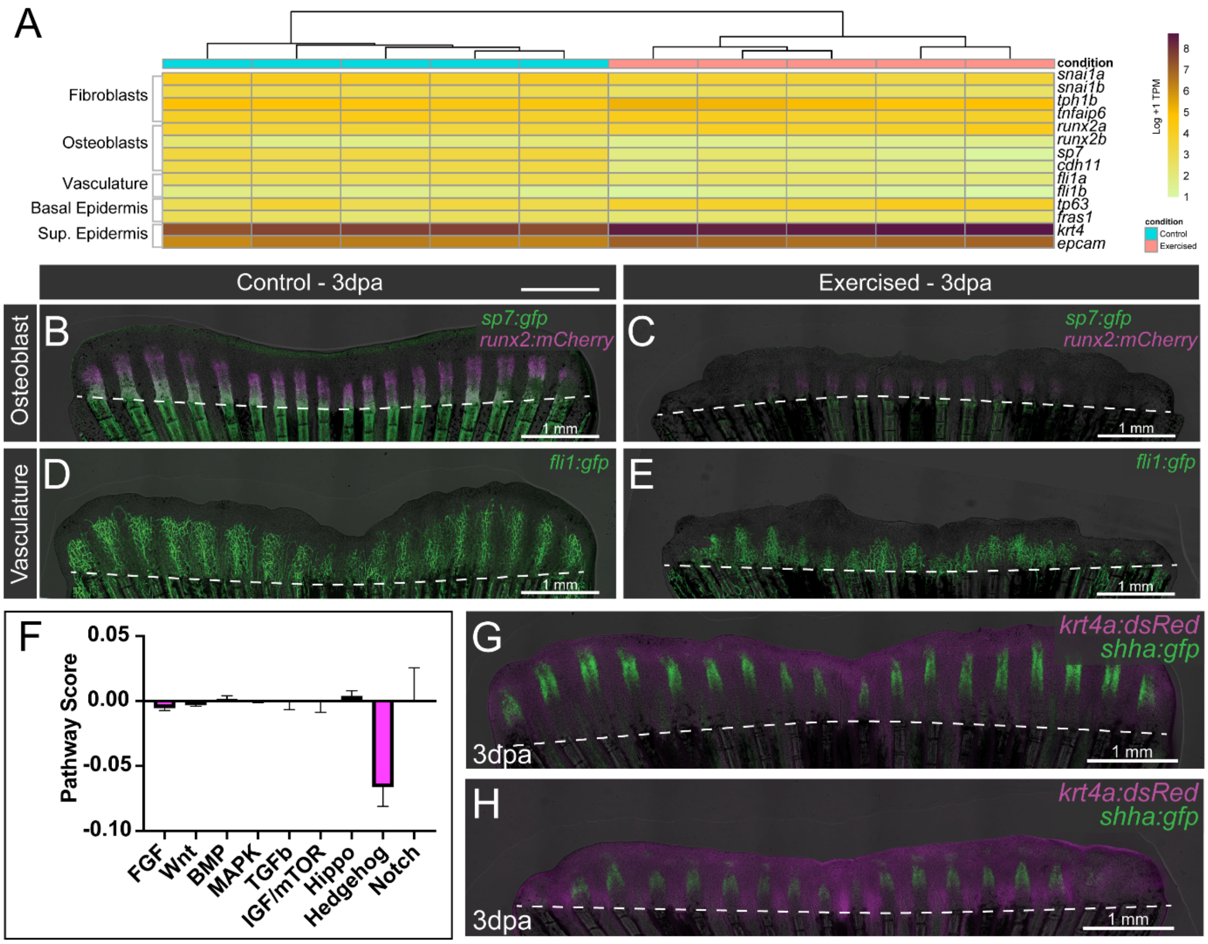


**Supplemental Figure 5. Exercise impacts on lineage-specific gene expression and common developmental signaling pathways during fin regeneration.** (A) Heatmap and hierarchical clustering of RNA-Seq replicates showing the expression levels of markers for fibroblasts (*snai1a/b, tph1b, tnfaip*), osteoblasts (*runx2a/b, sp7, cdh11*), vasculature (*fli1a*/*b*), basal epidermis (*tp63, fras1*) and superficial epidermis (*krt4, epcam*) in control and exercised conditions. (B, C) Representative images at 3 dpa showing Tg(*sp7:EGFP*) osteoblast and Tg(*runx2:mCherry*) pre-osteoblast lineage reporter expression and (D, E) 3 dpa Tg(*fli1:gfp*) vascular lineage reporter expression in control (B, D) and exercised (C, E) conditions following 3 days of swimming exercise at 50% maximum swim speed (*n*=6 fish per group). (F) Pathway score (Z-score) analysis of RNA-Seq fold change of common signaling pathway components (FGF, Wnt, BMP, MAPK, TGFβ, IGF/mTor, Hippo, Hh and Notch). (G, H) Representative images showing 3 dpa expression of Tg(*shha:gfp*) Sonic Hedgehog and Tg(*krt4a:mCherry*) superficial epidermal lineage reporters in control (G) and exercised (H) conditions following 3 days of swimming exercise at 50% maximum swim speed (*n*=6 fish per group). Dashed white lines indicate the amputation plane. Scale bars represent 1 mm. Error bars show standard deviation.

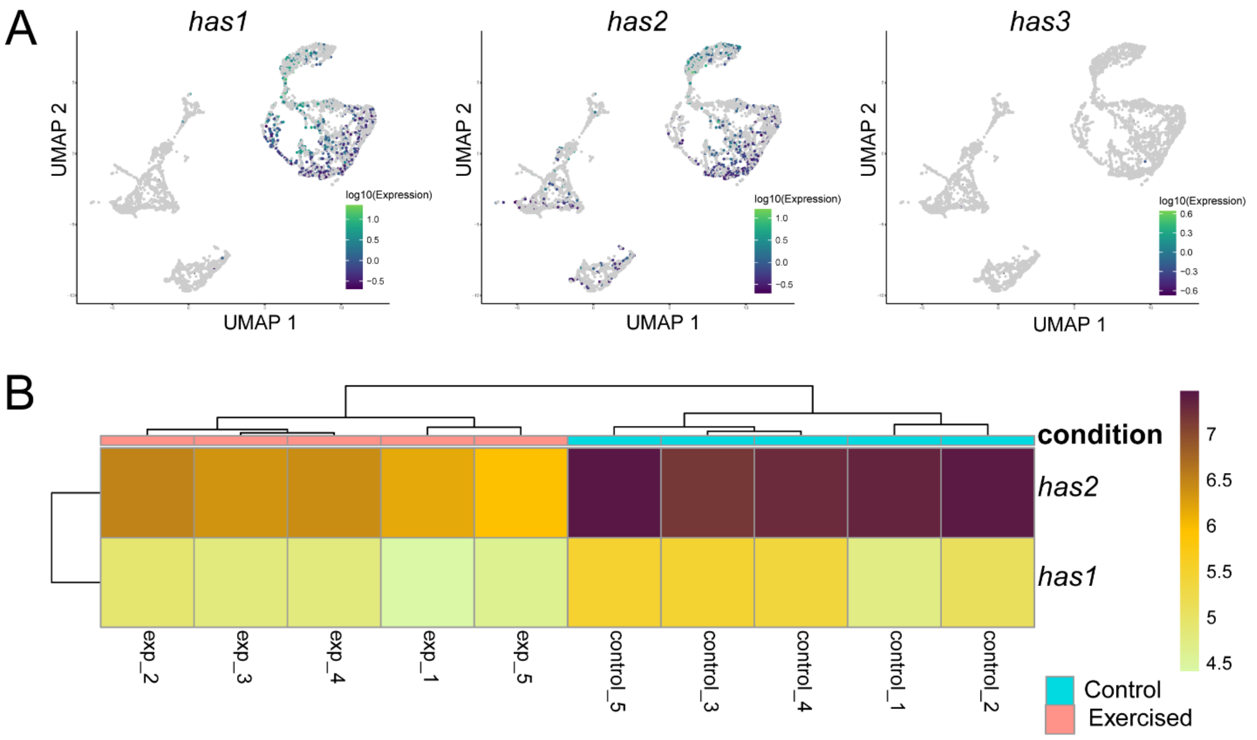


**Supplemental Figure 6. *hyaluronic acid synthase* (*has*) genes are enriched in fibroblast/osteoblast lineage cells and suppressed by exercise during fin regeneration.** (A) UMAP plots showing the expression of *has1*, *has2*, and *has3* genes from a combined 3 and 7 dpa scRNA-seq dataset. (B) Heatmap and hierarchical clustering of RNA-Seq replicates showing the expression levels of *has1* and *has2* transcripts under control and exercised conditions.

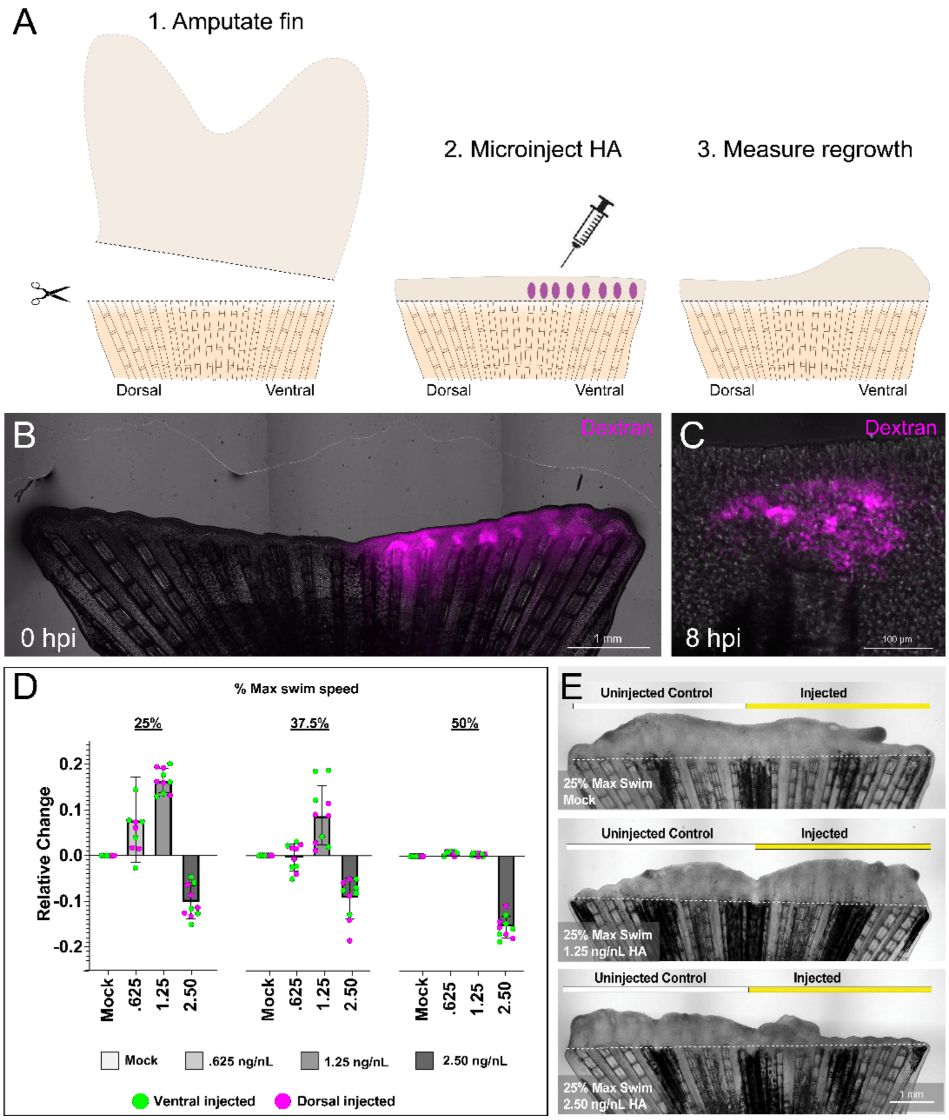


**Supplemental Figure 7. Exogenous HA prior to swimming ameliorates deleterious exercise effects.** (A) Cartoon schematic detailing HA intra-blastemal experimental procedure. (B) Representative whole mount 3 dpa regenerating caudal fin at 0 hours post-injection (hpi) with fluorescent dextran (magenta) overlay. (C) Representative area of a whole mount 3 dpa regenerating caudal fin blastema at 8 hpi with fluorescent dextran (magenta) overlay. (D) Graphs showing the change in regenerative area relative to uninjected controls following intra-blastemal injection of exogenous HA (*n*= 5 animals/trial and 2 replicate trials). 10 nL solutions of 0.625 ng/nL, 1.25 ng/nL, or 2.50 ng/nL HA were injected into each blastema prior to swim exercise at 25%, 37.5%, and 50% of maximum swim speed. Unnormalized data points were calculated by uninjected area / injected area. (E) Representative images of uninjected control, 1.25 ng/nL HA and 2.50 ng/nL HA intra-blastemal injected caudal fins after two swim exercise sessions at 25% maximum swim speed. Dashed white lines indicate the amputation plane. Scale bars in B and E represent 1 mm; C is 100 µm. Error bars are standard deviations.

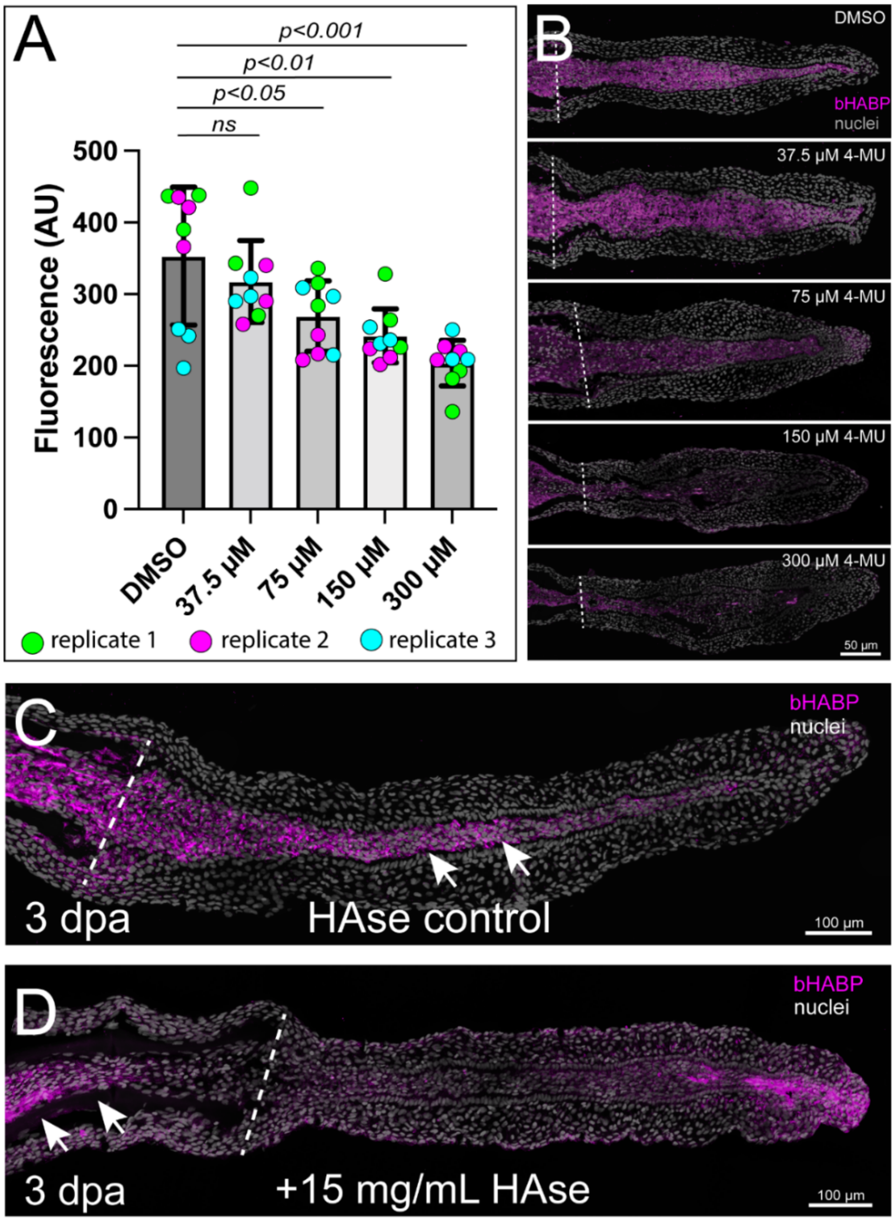


**Supplemental Figure 8. 4-methylumbelliferone or hyaluronidase treatment depletes hyaluronic acid during regeneration. (A)** Quantification of bHABP fluorescence intensity (AU) indicating HA levels in fin tissues treated with DMSO or indicated concentrations of small molecule HA-inhibitor 4-MU (*n*= 3 sections/replicate and 3 replicate animals). **(B)** Representative images of regenerating caudal fin transverse sections at 3 dpa showing bHABP staining (magenta) after treatment with DMSO or 4-MU at concentrations shown in A. **(C, D)** Representative images of regenerating fin transverse sections at 3 dpa showing bHABP staining (magenta) after either mock blastemal injections (C) or intra-blastemal injection of 15 mg/mL HAse. Arrows indicate bHABP staining in the blastema of mock injected (C) or intra-ray fibroblasts proximal to the amputation plane of HAse injected animals. Dashed white lines indicate the amputation plane. Scale bars represent 50 μm in panel B and 100 μm in panels C and D. Significance determined by one-way ANOVA with Dunnett's post-hoc test. Error bars represent standard deviation.

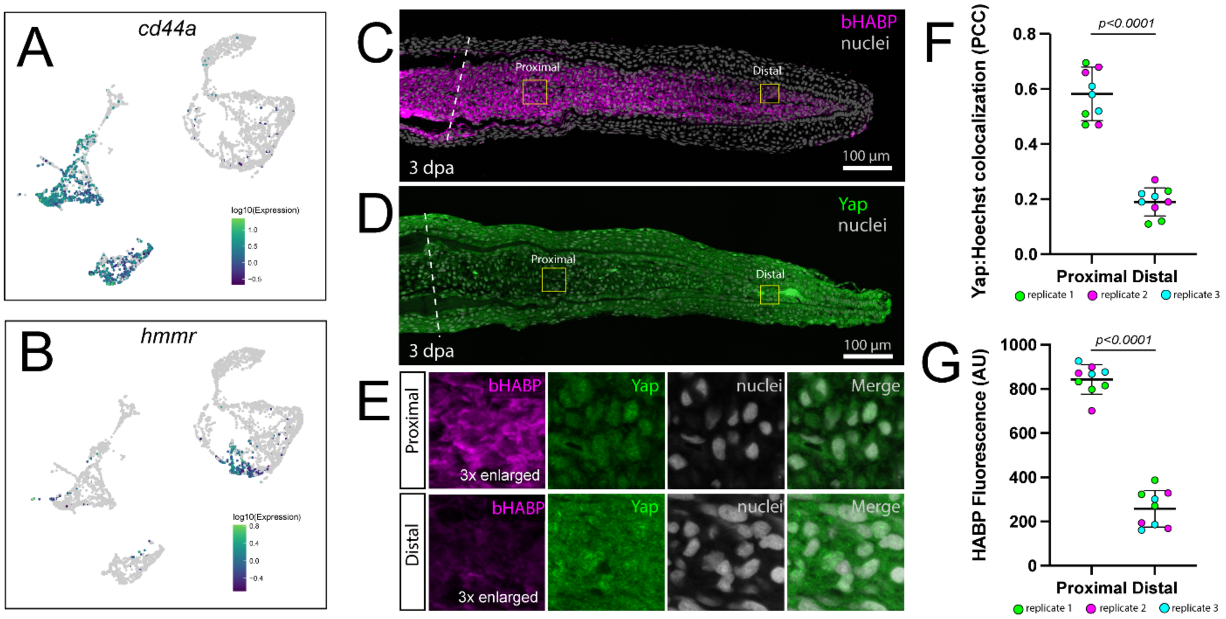


**Supplemental Figure 9. bHABP staining intensity correlates with Yes-associated protein (Yap) nuclear localization.** (A, B) UMAP plots showing the lineage-specific expression of canonical HA-ligand receptor/effector *cd44a* (A) and *hmmr* (B), respectively, from a subset combined 3 and 7 dpa scRNA-Seq dataset. (C) Representative image of a regenerating fin section at 3 days post-amputation (dpa) stained for bHABP (magenta). (D) Antibody staining of 3 dpa section for Yap (green). (E) Enlarged views of the boxed areas in C and D, showing expression of bHABP and Yap nuclear localization in proximal and distal regions. (F) Quantification of Yap-Hoechst colocalization (Pearson correlation coefficient, PCC) in proximal and distal regions in 3 dpa regenerating fin sections (*n*=3 sections/replicate and 3 replicate animals). (G) Quantification of HA binding protein (HABP) fluorescence intensity (AU) in proximal and distal regions in 3 dpa regenerating fin sections (*n*=3 sections/replicate and 3 replicate animals). Dashed white lines indicate the amputation plane. Scale bars represent 100 µm. Significance determined by paired two-tailed Student's t-tests. Error bars show standard deviation.

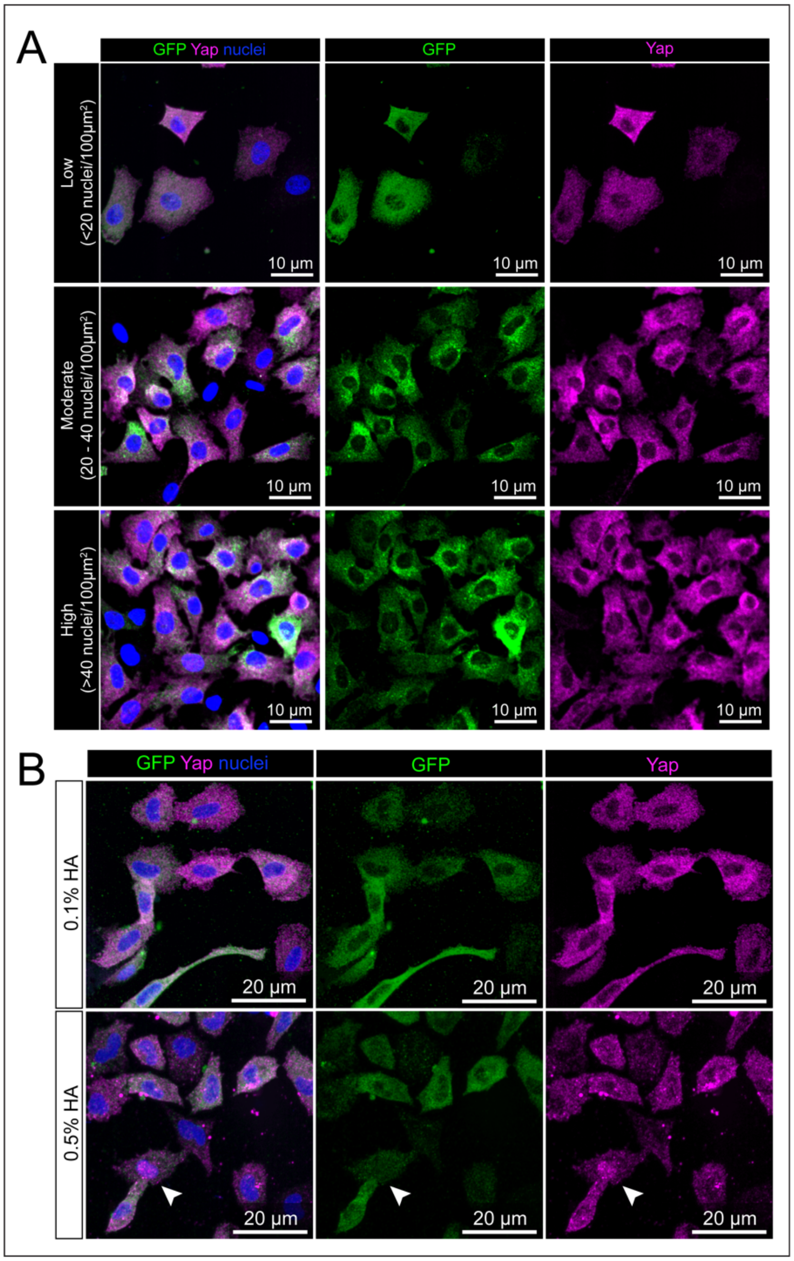


**Supplemental Figure 10. Yap nuclear localization in primary fin fibroblasts is independent of randomly seeded cell densities. (**A) Representative immunofluorescence images of primary fin blastema-derived fibroblasts cultured overnight on collagen-coated coverslips at varying cell densities: low (<20 nuclei/100 μm$^2$), moderate (20–40 nuclei/100 μm$^2$), and high (>40 nuclei/100 μm$^2$) ($n$=3). Cells were stained for GFP (green; marking *tph1b:GCaMP6s*$^+$ fibroblasts), Yap (magenta), and nuclei (blue; Hoechst). Yap nuclear localization appeared low at all cell densities. (B) Representative images of primary fibroblasts cultured on 0.1% collagen supplemented with either 0.1% or 0.5% hyaluronic acid (HA) ($n$=9 over 3 replicate trials). Cells treated with 0.5% HA show slightly enhanced but insignificant nuclear Yap accumulation (arrowheads). Scale bars as indicated.

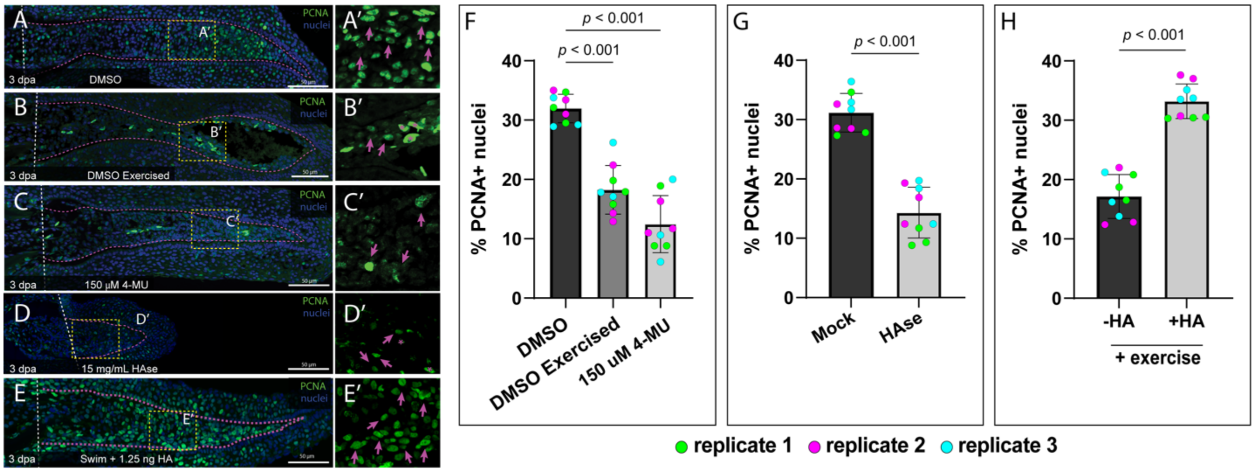


**Supplemental Figure 11. Proliferation in regenerating caudal fins after HA depletion.** (A–E) Representative images of 3 dpa regenerating caudal fin sections stained for Proliferative Cell Nuclear Antigen (PCNA; green) and nuclei under different conditions: (A, A') Unexercised with DMSO. (B, B') Swim exercised. (C, C') 3 day treatment with 150 µM 4-methylumbelliferone (4-MU). (D, D') Blastemal microinjection of 15 mg/mL HAse. (E, E') Swim exercised following blastemal microinjection of 1.25 ng HA. The boxed areas in panels A–E are shown enlarged in panels A'–E' to highlight PCNA-positive nuclei (magenta arrows). (F-H) Quantification of PCNA-positive nuclei following (F) DMSO, exercised or 150 µM 4-MU treatment, (G) mock or intra-blastemal injection of 15 ng/nL HAse, and (H) with and without intra-blastemal injection of 1.25 ng/nL HA prior to exercise ($n$= 3 sections/replicate and 3 replicate animals). Dashed white lines indicate amputation plane. Scale bars represent 50 µm. Significance in (F) determined by one-way ANOVA with Dunnett's post-hoc test. Significance in (G, H) determined by paired two-tailed Student’s t-tests. Error bars show standard deviation.
